# MerCat2: a versatile *k*-mer counter and diversity estimator for database-independent property analysis obtained from omics data

**DOI:** 10.1101/2022.11.22.517562

**Authors:** Jose L. Figueroa, Ajay Panyala, Sean Colby, Maren Friesen, Lisa Tiemann, Richard Allen White

## Abstract

**Summary:** MerCat2 (“<u>Mer</u> - <u>Cat</u>enate<u>2</u>”) is a versatile, parallel, scalable and modular property software package for robustly analyzing features in omics data. Using massively parallel sequencing raw reads, assembled contigs, and protein sequences from any platform as input, MerCat2 performs *k*-mer counting of any length *k*, resulting in feature abundance counts tables, quality control reports, protein feature metrics, ecological diversity metrics, and graphical representation (i.e., PCA). MerCat2 allows for direct analysis of data properties in a database-independent manner that initializes all data, which other profilers and assembly-based methods cannot perform. MerCat2 represents an integrated tool to illuminate omics data within a sample for rapid cross-examination and comparisons.

**Availability and implementation:** MerCat2 is written in Python and distributed under a BSD-3 license. The source code of MerCat2 is freely available at https://github.com/raw-lab/mercat2. MerCat2 is compatible with Python 3 on Mac OS X and Linux. MerCat2 can also be easily installed using bioconda: conda install MerCat2.

**Contact:** Richard Allen White III, UNC Charlotte

**Supplementary information:** Supplementary data are available online.

## 1 Introduction

Massively parallel sequencing (MPS) of whole ecosystems has elucidated the complexity of microbial composition, functions, and potential roles. With the scaling of terabytes of sequencing data, the illumination of genomes is ever more present, resulting in the expansion of reference databases needed for matching data using compositional, functional, and assembly-based methods. Kraken2 (Wood *et al*., 2019), MetaPhlAn2 (Truong *et al*., 2015), DRAM (Shaffer *et al*., 2020), and MicrobeAnnotator (Ruiz-Perez *et al*., 2021) provide database-dependent analysis of composition and function directly from metaome data (e.g., metagenomics and/or metatranscriptomics). However, database-dependence approaches fail to utilize, classify, or model all data due to a lack of references within a database. In addition, *de novo* assembly approaches, although recently improved for complex data types (Howe *et al*., 2014; Li *et al*., 2015), cannot assemble all data into contigs, leaving data underutilized. Open-source reference sequence databases are facing several challenges, including finding ongoing funding; many are moving to subscription-based access (e.g., KEGG www.kegg.jp/kegg/) and discontinuation (e.g., CAMERA http://camera.calit2.net/). Therefore, robust tools are needed to analyze these data in a database-independent manner.

Database-independent property analysis (i.e., DIPA), which utilizes counting of *k*-mer subsequences (of length *k*) from sequence reads without a reference sequence database for matching query data. DIPA-based *k*-mer counting provides rapid and robust microbial community analysis and characterization without the biases or limitations of sequence databases (Jiang *et al*., 2012) and/or *de novo* assembly to compare and contrast sequence datasets. *K*-mers are critical to assembly (Li *et al*., 2015), counting (Zhang *et al*., 2014), partitioning (Howe *et al*., 2014), genomic binning (Wu *et al*., 2015), and classification (Jiang *et al*., 2012). *K*-mer-based counting is amongst the fastest approaches for profiling metaomic data (Lindgreen *et al*., 2015). MerCat2 improves on MerCat v1 (White III *et al*., 2017) with greater parallelization, more considerable scalability, and added visualizations.

Here we describe MerCat2, a tool that can accommodate any size sequence file by utilizing a ‘divide and conquer’ approach that performs integrated analysis, including quality control, *k*-mer counting, and visualization. MerCat2 provides a rapid, robust, versatile analysis of MPS data using DIPA.

## 2 FEATURES

MerCat2 is a versatile and scalable Python-based open-source software package. For massive parallel processing (MPP) and scaling, we developed a byte chunking algorithm 1 (’Chunker,’ see Supplementary Figure S1) to split files for MPP and utilization in RAY, a massive open-source parallel computing framework to scale python applications and workflows (www.ray.io/). We also implemented a naive *k*-mer counter algorithm in base python for a range of sizes of *k* for versatile counting of both nucleotide and amino acid fasta files (see Supplementary Figure S2). However, merging large *k*-mer count tables as tabular delimited files (as .tsv) required large memory consumption (>50 GB of RAM). The Dask library we executed in our previous MerCat v1 was responsible for this extensive RAM utilization. To avoid large RAM consumption for streamlining upon low memory systems (e.g., a laptop), we implemented a greedy algorithm that limits RAM usage in native Python even for large datasets (>100 GB of raw sequence data and >60,000 bacterial genomes) (see Supplementary Figure S3). To plot large PCAs, which was not a previous feature of MerCat v1, we have utilized and modified an incremental PCA function from sci-kit learn (see Supplementary Figure S4). MerCat2 to scales from laptop to high-performance computing resources, all within the same user-friendly package.

MerCat2 computes *k*-mer frequency counting to any length *k* on assembled contigs as nucleotide fasta, raw reads or trimmed (e.g., fastq), and translated protein-coding open reading frames (ORFs) as a protein fasta (i.e., fasta amino acid file, .faa). The package also allows for user-defined custom analyses. Although raw read inputs can be used in MerCat2, it is not recommended due to low quality and sequencing errors. Instead, we utilize fastp (Chen *et al*., 2018) for quality control trimming of low-quality data obtained by fastq formats (default trimming is base pair quality score >Q_30_). For raw reads, MerCat2 provides fastqc reports pre- and post-trimming, which are also included within the final HTML-based report (Andrews, 2010). Outputs include tabular *k*-mer frequency count tables, tabular ecological diversity metrics (e.g., Alpha diversity), compositional and property dashboard, and PCA ordination (>4 samples). In addition, Alpha diversity metrics chao1, ACE, Simpson, Good’s coverage, dominance, and Fisher’s are provided in tabular format.

MerCat2 has two analysis modes utilizing nucleotide or protein files. In nucleotide mode, outputs include %G+C and %A+T content, contig assembly statistics, and raw/trim read quality reports are a provided output. For protein mode, nucleotide files (i.e., reads and contigs) can be translated into ORFs using Prodigal (Hyatt *et al*., 2012), which is prokaryotic specific or FragGeneScanRs (Van der Jeugt *et al*., 2022), which works well for general ORF calling. FragGeneScanR provides better gene calling for eukaryotic-rich samples than highly prokaryotic-rich samples. Prodigal is explicitly for prokaryotic gene calling due to the utilization of the Shine-Dalgarno sequence identification (i.e., a ribosomal binding site sequence) that is preferentially found in bacteria, and some archaea but not in eukaryotes (Hyatt *et al*., 2012). Protein property metrics of translated ORFs are provided in tabular format for protein isoelectric point (pI) and hydrophobicity metrics.

## 3 USE CASES

To demonstrate the scaling, versatility, and robustness of MerCat2, we compared large microbial genome databases, metagenomic and metatranscriptomic data from a diversity of samples.

### Five genomes example data set

Our general test data set includes five bacterial genomes ranging in GC/AT content with high GC >75% *Agrococcus pavilionensis* strain RW1 (White III *et al*., 2018) and the rest with moderate GC content 50-61% *Azotobacter vinelandii* strain DJ, *Rhizobium leguminosarum*, and two genomes of *Exiguobacterium chiriqhucha* strain RW2 and strain GIC31 that are highly similar (i.e., strains of each other) >97% using average nucleotide identity (White III *et al*., 2019). For both 4- and 31-mers, MerCat2 can complete counting in <6 secs, <12 GB of RAM, and <500 Mb of disk usage (**Table 1**).

**Table 1:** Use case data statistics. Time in seconds, RAM in Gigabytes, and Disk space used as Kilobytes to MegaBytes. Counting tests uses a minimum *k*-mer count of 10 by default settings.

| Dataset | kmers | Time (S) | RAM (GB) | Disk Usage |
| --- | --- | --- | --- | --- |
| Archaea AA | 4 | 23.08 | 12.58 | 699M |
| Archaea AA | 31 | 28.44 | 12.97 | 352K |
| Archaea NT | 4 | 29.72 | 11.66 | 11M |
| Archaea NT | 31 | 90.92 | 8.96 | 32M |
| Bacteria AA | 4 | 715.87 | 14.91 | 20G |
| Bacteria AA | 31 | 801.59 | 11.87 | 841M |
| Bacteria NT | 4 | 999.36 | 27.7 | 389M |
| Bacteria NT | 31 | 3019.21 | 27.58 | 9.9G |
| Soil AA | 4 | 889.99 | 32.14 | 9.4M |
| Soil AA | 31 | 699.33 | 31.35 | 3.1M |
| Soil NT | 4 | 2663.12 | 24.2 | 40K |
| Soil NT | 31 | 5056.6 | 19.35 | 28M |
| SharkBay AA | 4 | 16.64 | 18.01 | 6.6M |
| SharkBay AA | 31 | 9.43 | 23.8 | 32M |
| SharkBay NT | 4 | 53.49 | 12.48 | 36K |
| SharkBay NT | 31 | 101.02 | 15.53 | 228M |
| Test 5 genomes NT | 4 | 2.07 | 2.47 | 8K |
| Test 5 genomes AA | 4 | 0.96 | 3.84 | 1.2M |
| Test 5 genomes NT | 31 | 5.59 | 2.48 | 2.4K |
| Test 5 genomes AA | 31 | 1.24 | 3.51 | 12K |

### GTDB archaea/bacterial

We used the GTDB archaea genome database (3,412 species) and GTDB bacterial genome (62,291 bacterial species) (Park et al., 2022) to test scalability, disc space, and memory use of *k*-mer counting and PCA plotting. This dataset represents one of the largest high-quality controlled collections of archaea and bacteria genomes. For the GTDB archaea genome database, MerCat2 was able to complete counting whether 4- or 31-mer as either nucleotides or amino acids in under <100 secs, using <15 GB of RAM and <700 Mb of disc space (**Table 1**). MerCat2 counted GTDB bacterial genome database with whether 4- or 31-mer as either nucleotides or amino acids in <1 h, <30 GB of RAM, and <10 GB of disc space (**Table 1**).

### Shark Bay Metatranscriptomes

We included metatranscriptome samples from modern hypersaline microbial mats to test scalability, disc space, and memory for MerCat2. These are ten samples with five replicate each from smooth and pustular mats from the Nilemah tidal flat located in the southern area of Hamelin Pool, Shark Bay, Western Australia (Campbell *et al*., 2019). The data for all the Shark Bay metatranscriptomes was 15 GB. All ten samples were counted with either 4- or 31-mer as either nucleotides or amino acids in under 2 mins, <20 GB of RAM, and <250 Mb of disc space (**Table 1**).

### Switchgrass soil metagenomes

Eight large soil metagenomes isolated from Lux Arbor, Michigan, were used to test *k*-mer counting and PCA plotting scalability, disc space, and memory use. The combined datasets are >5 billion reads, ~671 million per sample, at >100 GB of data per sample. Scalability for 5 billion metagenomic reads was possible with MerCat2 as it counted all eight datasets with whether 4- or 31-mer as either nucleotides or amino acids in under 90 mins, <35 GB of RAM, and <10 Mb of disc space (**Table 1**).

## 4 CONCLUSIONS

MerCat2 provides DIPA for metaomic data, utilizing all data present, in a robust, versatile manner resulting in tabular files for downstream analysis, with built-in visualizations and a dashboard. MerCat2 is scalable, accommodating for various input files, user-friendly, easy to install, and user-customizable. MerCat2 enables rapid analysis of many datasets and large datasets in a database-independent manner.

**Figure 1:**
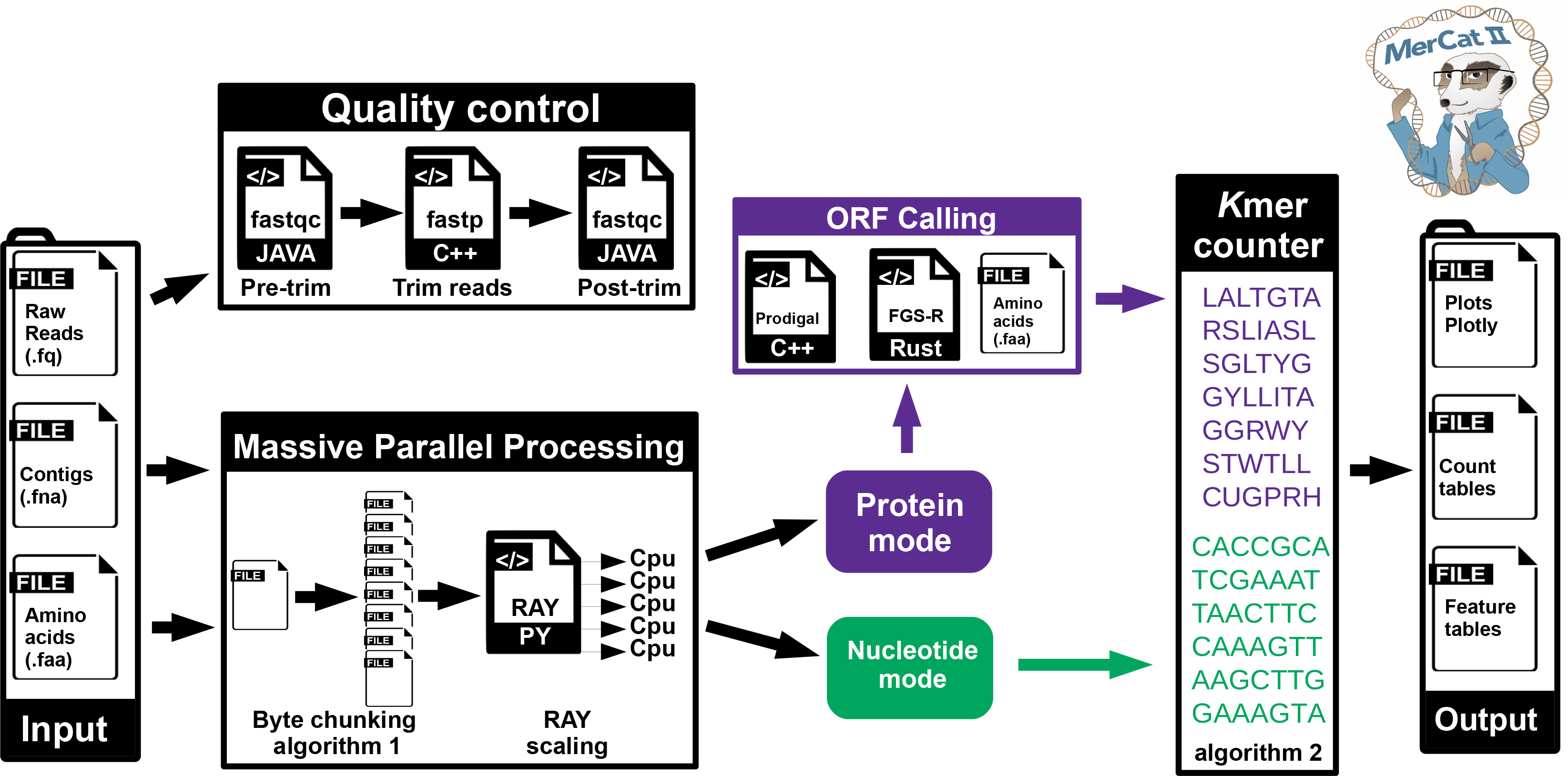
Flow graph of MerCat2. MerCat2 computes *k*-mer frequency counting to any length *k* on assembled contigs as nucleotide fasta, raw reads or trimmed (e.g., fastq), and translated protein-coding open reading frames (ORFs) as a protein fasta (i.e., fasta amino acid file, .faa) as inputs. On raw reads, it performs quality control using fastqc (pre/post) trimming with fastp. The massive parallel processing (MPP) occurs with an integration of the byte chunking algorithm 1 (Chunker - Supplemental Figure S1) with the python library RAY for scaling. MerCat2 can be run in protein or nucleotide mode using the naive *k*-mer counter algorithm 2 (*k*-mer counter - Supplemental Figure S2). For protein mode, the user can choose Prodigal or FragGeneScanR to convert to ORFs. No ORF calling is required if the user already has an amino acid fasta. Outputs include *k*-mer frequency count tables, tabular ecological diversity metrics (e.g., Alpha diversity metrics), protein and nucleotide feature tables, compositional and property dashboard, and PCA ordination (>4 samples).

## Acknowledgments

We thank Kurt R Glaesemann, Kevin Glass, Janet, and Christer Jansson for supporting the original MerCat v1 at PNNL. Nathan Johnson for his assistance in preparing excellent figures.

## Data Availability

The data underlying this article are available at github.com/raw-lab/mercat2 and https://osf.io/mzrvj/. Shark Bay metatranscriptomes are in OSF under “Shark bay Transcriptomes” at https://osf.io/e94yg/. Lux Arbor soil metagenomes are in OSF under “MMPRNT panicum metagenome mags” at https://osf.io/mzrvj/. For GTDB data is available at https://data.gtdb.ecogenomic.org/releases/release207/

## Supplemental Figure Legends

**S1**. Byte chunking algorithm 1 description (Chunker)

**S2**. Naive *k*-mer counter algorithm 2 description

**S3**. Tab-separated merging algorithm 3 description

**S4**. Incremental PCA algorithm 4 description modifications

