## Supplemental Figures for "MerCat2: a versatile *k*-mer counter and diversity estimator for database-independent property analysis obtained from omics data"

**Algorithm 1:** Chunker

```
Input  : Fasta file
Result : multiple smaller fasta files
count ← 1
newSequence ← empty string
foreach sequence in fasta_file do
    newSequence ← sequence
    sample ← list of chunks
    if newSequence > chunkSize then
        filename ← chunk + count
        write file filename
        newSequence ← empty string
    end
end
```

**Figure S1**

**Algorithm 2:** k-mer counter

```
Input  : List of sequence files, optionally split in smaller chunks
Result : TSV file with k-mer counts
Function kmer_counter(fasta_file):
    foreach sequence in file do
        counts ← kmer counts using sliding window
    end
    return counts

foreach file in files do
    foreach chunk in file do
        kmer_counter(chunk) → RAY for MPP
    end
    counts ← merged kmer counts
end
```

**Figure S2**

**Algorithm 3:** Tab Separated File merger

```
Input  : List of sequence files
Result : TSV file with k-mer counts

names ← List of sample names
out_file ← path to combined tsv file
out_file ← write header with sample names
foreach sample in names do
    open all tsv files
end
kmer_set ← empty set
foreach sample in names do
    kmer_set ← add k-mer name
    curr_line[sample] ← first kmer & count in sample TSV
end
cur_kmer ← first kmer fromkmer_set
while True do
    next_line ← empty list
    foreach sample in names do
        if cur_line[sample][kmer] > cur_kmer then
            next_line ← append 0
        else
            next_line ← append cur_line[sample][kmer count]
            curr_line[sample] ← next kmer & count in sample TSV
            kmer_set ← add k-mer name
        end
    end
    write next_line to out_file
    if kmer_set is empty then
        break loop
    end
end
```

**Figure S3**

**Algorithm 4:** Incremental PCA

```
Input  : List of sequence files
Result : TSV file with k-mer counts
PCA ← import Incremental PCA from sklearn in Python
chunk_size ← 1000
dataframe ← file handle for reading in TSV
foreach chunk in dataframe do
    PCA.partial_fit(chunk)
end
foreach chunk in dataframe do
    PCA.transform(chunk)
    append data to output TSV
end
Plot 3D PCA with Plotly
```

**Figure S4**
